# Long-lasting Avoidance Induced by Repeated Stimulation of the Rostromedial Tegmental Nucleus

**DOI:** 10.1101/2025.05.30.657063

**Authors:** J.R. Watson, M.D. De Leon Lopez, E.L. Carlson, P.J. Vento

## Abstract

Adaptive decision-making requires the ability to weigh the relative costs versus the benefits of our actions and to flexibly respond to changing environmental contingencies. While midbrain dopamine (DA) has been a major focus of study for its role in associative learning, motivation, and reward seeking, far less is known regarding the contribution of inhibitory dopamine inputs in promoting behavioral inhibition and avoidance learning. The rostromedial tegmental nucleus (RMTg) sends dense GABAergic projections to DA neurons in the ventral tegmental area (VTA) and mounting evidence suggests the RMTg–VTA circuit is required to suppress reward seeking under punishment, but it remains unclear whether stimulation of this pathway is sufficient to drive learning and promote shifts in cost-benefit decisions. To test this, we developed two separate tasks of passive and active avoidance in rats where lever pressing for food reward was associated with repeated contingent optogenetic stimulation of the RMTg–VTA pathway. In a one-lever fixed ratio 5 food-seeking task, we found that pairing RMTg–VTA stimulation with reward delivery caused a robust, yet transient suppression of reward seeking that quickly returned to baseline after contingent stimulation ceased. When given an alternative reward choice, however, RMTg–VTA stimulation caused a rapid and enduring shift in subjective choice leading to persistent and selective avoidance of the stimulation-paired reward. These findings support a multifaceted role for the RMTg–VTA pathway in learning and decision-making and provide key insights into the neural mechanisms underlying behavioral avoidance and maladaptive reward seeking.

## 1.0 Introduction

Pursuing rewards while minimizing potential costs is a fundamental component of adaptive decision-making that guides behavior across species [1, 2]. Efficient decision-making, therefore, requires not only the motivation to seek out beneficial outcomes but also the flexibility to suppress ongoing (and future) reward-seeking behavior when negative consequences are incurred [3, 4]. The ability to flexibly modify behavior in response to suboptimal outcomes is commonly disrupted in a variety of psychiatric conditions including mania and substance use disorder, where individuals persistently seek out rewards despite often profound negative consequences [2, 5, 6]. A more complete understanding of the neural mechanisms underlying behavioral inhibition and adaptive decision-making is critical for identifying the neurobiological substrates of such compulsive disorders and how individuals learn to optimize positive outcomes while avoiding potential threats.

While substantial research has focused on the role of midbrain ventral tegmental area (VTA) dopamine (DA) neurons in reward seeking and reinforcement learning [2, 7–9], it is increasingly recognized that inhibitory inputs to the VTA and DA neuron inhibition likely play similarly important roles in aversive learning and avoidance [10]. For example, evidence shows that VTA DA neurons are inhibited by aversive outcomes, and that this inhibition plays a critical role in guiding behavioral adaptation [11–13]. Danjo et al. (2014) demonstrated that real-time optogenetic inhibition of VTA DA neurons in a conditioned place avoidance (CPA) paradigm produced an avoidance response that persisted beyond conditioning, suggesting that VTA DA inhibition may serve as a teaching signal to promote enduring avoidance responses.

The rostromedial tegmental nucleus (RMTg) is a GABAergic midbrain structure that exerts robust inhibitory control over VTA DA neurons [15, 16] with emerging roles in modulating reward seeking and aversive learning [17–20]. Notably, the RMTg is activated by a broad range of aversive stimuli, including footshock, shock-associated cues, reward omission, and drug withdrawal [16, 17, 21–23], and stimulation of RMTg axon terminals in the VTA produces real-time place aversion [24]. Conversely, inactivation of the RMTg–VTA pathway induces persistent reward seeking despite impending footshock punishment, suggesting that when RMTg signaling to the VTA is disrupted, aversive stimuli fail to suppress reward-seeking behavior, leading to punishment resistance [18]. Collectively, these findings support the hypothesis that the RMTg acts as a ‘GABAergic brake’ on DA signaling to inhibit behavioral responding in the presence of aversive outcomes [25, 26]. While the role of the RMTg in suppressing behavior in response to aversive environmental stimuli is coming into focus, much remains unknown regarding how causal activation of the RMTg–VTA pathway shapes decision-making across different environmental contexts and diverse behavioral contingencies.

In the present study we used operant learning tasks for natural (food) reinforcers in rats to test the causal effects of RMTg–VTA stimulation in avoidance learning. We found that when given a single response option in a one lever food-seeking task, contingent RMTg–VTA stimulation promotes behavioral inhibition in a manner similar to contingent footshock, suggesting a general suppressive influence of the RMTg in ongoing behavior. When rats are presented with multiple response options for distinct yet equivalent rewards, however, RMTg– VTA stimulation instead acts to promote active avoidance responses to bias subjective reward preferences away from choices associated with the stimulation, with this avoidance becoming more robust after repeated training and enduring long after stimulation ceased. Together, these data suggest the RMTg–VTA circuit drives distinct (and opposing) behavioral responses based on task contingencies to promote long-lasting changes in adaptive decision-making.

## 2.0 Methods

### 2.1 Animals

Adult male Sprague Dawley rats weighing approximately 300g upon arrival from vendor (Charles River Laboratories, Raleigh, NC) were individually housed in standard shoebox cages in a temperature-and humidity-controlled vivarium (12/12-hour light-dark cycle) with food and water provided *ad libitum*, unless stated otherwise. Experiments were performed during the light phase of the light/dark cycle. Approximately 3 days prior to behavioral testing, animals were food restricted to 85% ± 3% of their free-fed bodyweight. Thereafter, bodyweights were monitored and rats fed daily after behavioral testing. All procedures described here were approved by the University of South Carolina *Institutional Animal Care and Use Committee* and conformed to the National Institutes of Health *Guide for the Care and Use of Laboratory Animals*.

### 2.2 Stereotaxic Surgery

Under isoflurane anesthesia, rats were fixed in a stereotaxic frame, a small incision was made in the scalp, and small burr holes were drilled in the skull. Viruses were injected into the RMTg (10° angle; from bregma AP: -7.4mm, ML: +1.9mm; DV: -7.4mm from dura) and/or VTA (10° angle; from bregma AP: -6.0mm, ML: +2.3mm; DV: -8.5mm from skull) using Nanoject Auto-Nanoliter Injectors (100nL/min; Drummond Scientific Company, Broomall, PA) with pulled glass pipettes. Injectors were left in place for at least 5min post-infusion to permit diffusion of virus. For optogenetic studies, stainless-steel ferrules (2mm diameter, Doric Lenses, Franquet, Quebec, Canada) containing optical fibers were implanted bilaterally above RMTg axon terminals in the VTA (10° angle; from bregma AP: -5.4mm, ML: +2.3mm; DV: -7.8mm from skull). Implants were affixed to the skull using bone screws and dental acrylic. Ketoprofen (5mg/kg, sc) was administered before surgery and for up to two days thereafter, as needed, to reduce pain and swelling. At least 5 days were given for recovery prior to food restriction and/or behavioral training, and at least three weeks were allocated for viral transfection prior to testing neural manipulations.

### 2.3 Optogenetic Control of Neural Activity

Virus (UNC Vector Core, Chapel Hill, NC) encoding the light-sensitive cation channel channelrhodopsin-2 (AAV2-hSyn-hChR2(H134R)-mCherry) or control vector (AAV2-hsyn-mCherry) was injected bilaterally (400nL/hemisphere) into the RMTg and bilateral optical fibers were implanted in the VTA. Blue light (450nm) was delivered via computer-controlled laser (Dragon Lasers, Changchun, China; Laser Glow Technologies, North York, Ontario, Canada) mated to optical splitters by an optical rotary joint (Doric Lenses, Franquet, Quebec, Canada). Laser light (20mW/hemisphere) was pulsed at 33Hz (5ms duration).

### 2.4 Chemogenetic Control of Neural Activity

Retrogradely-transported virus encoding cre recombinase (pENN.AAVrg.hSyn.HI.eGFP-Cre.WPRE.SV40; Addgene, Watertown, MA) was injected into the VTA (150nL/hemisphere), and cre-dependent excitatory *designer receptors exclusively activated by designer drugs* (DREADDs; AAV-hSyn-DIO-hM3D(Gq)-mCherry; Addgene, Watertown, MA) or control vector (pAAV2-hSyn-DIO-mCherry; Addgene, Watertown, MA) was injected bilaterally into the RMTg (400nL/hemisphere). Twenty minutes prior to testing, the synthetic DREADDs ligand clozapine-N-oxide (CNO, 5mg/kg/ml in 8% DMSO) or vehicle (1ml/kg of 8% DMSO in sterile PBS) was injected intraperitoneally (IP).

### 2.5 Apparatus

Behavioral testing was conducted in standard operant chambers (30.5 X 24 X 21 cm; Med-Associates Inc.) enclosed in sound-attenuating boxes. Boxes were equipped with a ventilating fan that also served to obscure external noise. Operant chambers were arranged to either contain two retractable levers (one active, one inactive) with a cue light above each lever and a centralized food receptacle (Single Reward Task), or oriented with a centralized single retractable lever flanked on either side by two food pellet receptacles with a cue light above each receptacle (Dual Reward Choice Task). The wall opposite the lever(s) and receptacle(s) was equipped with a single house light. A photobeam was fitted in each food receptacle to monitor entries into the food port and to measure response latencies to the receptacle. Behavioral data were recorded on a computer using MedPC-V software connected to the operant chambers via an interface.

### 2.6 Single Reward Task

Rats were trained to lever press on a fixed ratio (FR) 5 for standard food pellets (45 mg; LabDiet, St. Louis, MO) in daily sessions comprising 70 trials or 60 minutes, whichever occurred first. Trials began with extension of both an active and inactive lever with cue lights illuminated above each lever. Completion of the FR5 resulted in delivery of a single food pellet, retraction of both levers, and the cue lights were extinguished. After a 15sec intertrial interval (ITI), a new trial was signaled by illumination of the house light and both levers extended. Rats completing >95% of trials in <15 seconds/trial for two consecutive sessions progressed to testing under one of three separate conditions. *Footshock*: Rats were tested in four once-daily sessions where completion of the FR5 resulted in delivery of a single food pellet accompanied by brief, mild footshock (500ms, 0.5mA) to evaluate the suppressive effect of a moderately aversive unconditioned stimulus on responding for food. *Optical stimulation*: Two additional groups of rats were tested in four once-daily sessions using the same task parameters as above, except instead of footshock, completion of the FR5 resulted in a single food pellet and optical RMTg– VTA stimulation. Rats received either 500ms stimulation, mimicking the duration of footshock in the first group, or 30sec stimulation to induce more robust activation of the RMTg–VTA circuit. Thereafter, rats in all three conditions (footshock, 500ms or 30sec stimulation) underwent three additional “recovery” sessions where footshock or optical stimulation were eliminated to test for lasting influences of the various manipulations on responding for food.

### 2.7 Dual Reward Choice Task

Rats were first trained to nosepoke in two opposing food pellet receptacles in an alternating fashion. Each trial was initiated by illumination of a cue light over the “active” receptacle, signaling reward availability on an FR1 schedule of reinforcement. After a successful nosepoke in the active receptacle, one of two potential food rewards was delivered: vanilla-or chocolate-flavored pellets, with location (left or right receptacle) counterbalanced between testing chambers. Once animals learned to reliably poke at each receptacle in <15sec on at least 95% of trials they progressed to the second phase of training. Phase two incorporated a central lever press (FR1) after nosepoking at the illuminated receptacle to receive the food reward. Once rats completed nosepoke and lever press in <15sec on at least 95% of trials over two consecutive sessions they advanced to free access choice sessions. Specifically, rats underwent three pre-stimulation choice sessions comprising 70 trials each where the cue lights above both receptacles were illuminated and rats were given the opportunity to freely respond for either reward option (vanilla or chocolate pellets) via nosepoke and lever press. For chemogenetics experiments, rats received IP injection of 0.9% saline (1ml/kg) immediately before the 3^rd^ and final pre-stimulation choice session to habituate them to injection, and for optogenetics experiments rats were connected to the optical splitter but no light was delivered for habituation to the optical cable. Baseline preference for each flavor was calculated as the average number of choices for the two reward options across the three pre-stimulation choice sessions, and the flavor chosen more frequently was designated the “preferred” flavor and paired with either CNO (chemogenetics) or light delivery (optogenetics). The “non-preferred” flavor was then paired with either vehicle (chemogenetics) or the optical splitter was attached but no light delivered (optogenetics). Chemogenetic stimulation sessions were designed to condition an avoidance to the preferred flavor through four CNO-paired sessions where rats had access to only the preferred flavor and four vehicle-paired sessions with access only to the non-preferred flavor (Figure 1A). Injections (CNO or vehicle) were performed on alternating days in a within-subjects design with at least 1 day between injections for drug washout. Optogenetic stimulation was performed over four consecutive sessions where responding for the previously identified “preferred” flavor resulted in 30sec of optical RMTg–VTA stimulation, while responding for the “non-preferred” flavor was accompanied by 30sec of no light delivery (Figure 1B). After stimulation sessions, rats underwent 10 additional post-stimulation “recovery” sessions to assess enduring effects of the previous RMTg–VTA stimulation.

**Figure 1:**
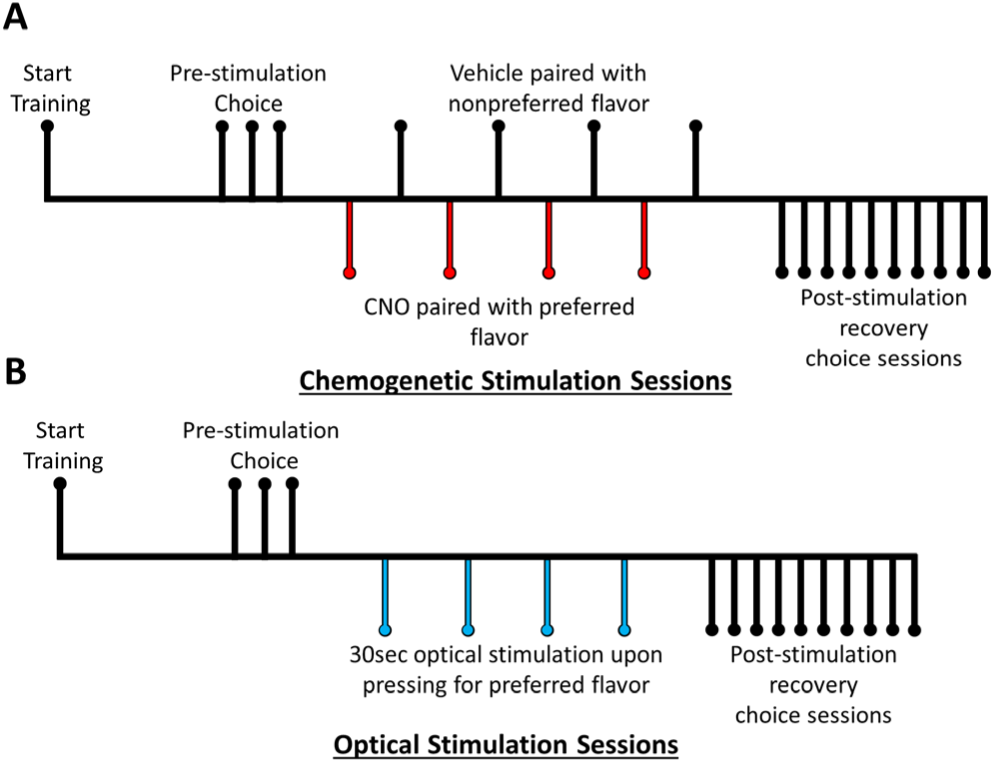
Timeline of the Dual Reward Choice Tasks. **A)** Schematic illustrating the experimental design of the chemogenetic approach including 3 pre-stimulation choice sessions, 8 alternating CNO-and vehicle-paired conditioning sessions, and 10 post-stimulation recovery “choice” sessions. **B)** Schematic illustrating the experimental design of the optogenetic approach which has replaced conditioning sessions with 4 light-paired real-time optical stimulation sessions.

### 2.8 Progressive Ratio Testing

Rats expressing ChR2 or mCherry control in RMTg with optical fibers implanted above RMTg terminals in the VTA underwent training and testing for the Dual Reward Choice Task using optogenetic stimulation, as described above. After the 4^th^ and final light-paired session, the following day rats were tested in twice-daily sessions over 4 consecutive days on a progressive ratio (PR) schedule of reinforcement for each reward (vanilla or chocolate). Each PR session included access to only one of the two reward options in an alternating, counterbalanced design, where the response requirement to receive a single vanilla or chocolate pellet progressively increased after each trial (e.g., 1, 2, 4, 9, 12, 15, 20, 25, 32, 40, 50, 62, 77, 95, 118, 145, etc.). Sessions continued until no response (lever press) was recorded for two consecutive minutes. The PR breakpoint was defined as the highest ratio completed before the session terminated due to inactivity.

### 2.9 Sucrose Preference Testing

After PR testing, *ad libitum* food access was restored, and the following day the same rats began sucrose preference testing (SPT). On the day of testing, food and water were removed from the home cage 15min prior to testing. Two bottles, one containing 5% sucrose solution and the other containing tap water, were then placed in the home cage in a counterbalanced orientation to control for potential side biases. After 1hr of free access to both solutions the bottles were removed and food/water were returned. Intake of each solution was assessed as the difference in bottle weight before/after testing. SPT occurred over 4 consecutive once-daily sessions.

### 2.10 Histology

Subjects were deeply anesthetized by inhaled isoflurane and transcardially perfused with 0.9% saline (150-200mL) followed by 10% formalin (∼150mL). Brains were removed and stored in 10% formalin overnight before transferring to 20% sucrose with 0.05% sodium azide for cryoprotection. Tissue was collected in 40µm-thick sections on a cryostat, and floating sections were processed using immunohistochemistry to verify viral expression and optical fiber placement. Tissue was incubated overnight in rabbit anti-tyrosine hydroxylase (TH, 1:1K, Abcam) or rabbit anti-FoxP1 (1:1K, Abcam) in phosphate buffered saline (PBS) with sodium azide (0.01%) and 0.25% Triton X-100 (Sigma-Aldrich). The transcription factor, FoxP1, has been established as a valid and reliable neuroanatomical marker of the RMTg [24] while TH was used to delineate the VTA. Fluorescence was visualized using secondary incubation in Alexa Fluor 488-conjugated donkey anti-rabbit (1:1000, Jackson Immunoresearch) in PBS for 30min. After each incubation, tissue was rinsed 3X in PBS at 1min/wash. Images were acquired on a DM6 Leica microscope.

### 2.11 Statistical Analyses

Repeated measures ANOVAs were conducted to evaluate differences in lever pressing, response latency, PR breakpoint, fluid intake, or the proportion of responses for preferred vs non-preferred rewards. Student’s T-tests were used to assess differences between the ChR2 and control groups in total fluid consumed in SPT. Bonferroni corrections were used to account for multiple comparisons. Greenhouse-Geisser corrections were used when sphericity assumptions for repeated measures ANOVAs were violated. Statistically significant main or interaction effects (p<0.05) were further probed using Student’s t-test.

## 3.0 Results

### 3.1 Contingent RMTg–VTA stimulation suppresses reward seeking under limited choice

While previous research has demonstrated that inhibition of the RMTg or its projections to the VTA profoundly impairs behavioral inhibition [17, 22, 27, 28], it has not yet been directly tested whether stimulation of the RMTg–VTA circuit is sufficient to suppress motivated behavior. To address this, rats expressing ChR2 or control virus in the RMTg with optical fibers implanted above the VTA (Figure 2A) were trained to lever press for food reward in the Single Reward Task (described above) where responding (FR5) was immediately followed by optogenetic RMTg–VTA stimulation. A separate group of rats were tested with response-contingent footshock (500ms, 0.5mA) to compare the suppressive effect of a moderately aversive unconditioned stimulus. Rats in the footshock condition robustly suppressed lever pressing for food beginning with the first exposure (Exp1) session and persisting across all 4 shock sessions (Exp1-Exp4). This was followed by a gradual resumption of lever pressing across recovery sessions (R1-R3) in the absence of shock (Figure 2B). Given that the RMTg is robustly activated by footshock, we next sought to mimic the suppressive effect of shock by optogenetically stimulating the RMTg–VTA pathway at the same time when footshock would have otherwise occurred, for the same duration of time (500ms), and at a frequency in the reported range of RMTg activation to aversive stimuli (33Hz) [17, 22]. Contingent 500ms optical stimulation did not, however, significantly alter responding as rats expressing ChR2 completed all trials during light sessions and recovery sessions (Figure 2B). Since 500ms stimulation was not effective at suppressing operant responding for food, we then increased the stimulation duration to 30 seconds per trial in a separate group of rats to evaluate whether more robust sustained activation of the circuit would model the behavioral suppression induced by footshock punishment. Indeed, in response to 30sec optical RMTg–VTA stimulation, rats expressing ChR2 exhibited robust suppression of lever pressing for food reward while the same duration of light exposure did not alter behavior in mCherry controls (main effect of Group (Shock, ChR2 500ms, ChR2 30sec, mCherry 30sec): *F*(3,23) = 53.495, *p* < 0.001; main effect of Session: *F*(2.478,56.984) = 33.316, *p* < 0.001; Group X Session: *F*(7.433,56.984) = 12.666, *p* < 0.001; Figure 2B). Notably, the suppressive effect of 30sec optical RMTg-VTA stimulation was not as strong as footshock (p<0.05, Exp1-Exp4), and neural stimulation was more transient compared to footshock punishment, as removal of the 30sec optical stimulation led to a rapid resumption of lever pressing in the first recovery session (R1) while rats in the footshock condition continued to significantly suppress responding at this time point (p<0.05 Shock vs all other groups).

**Figure 2:**
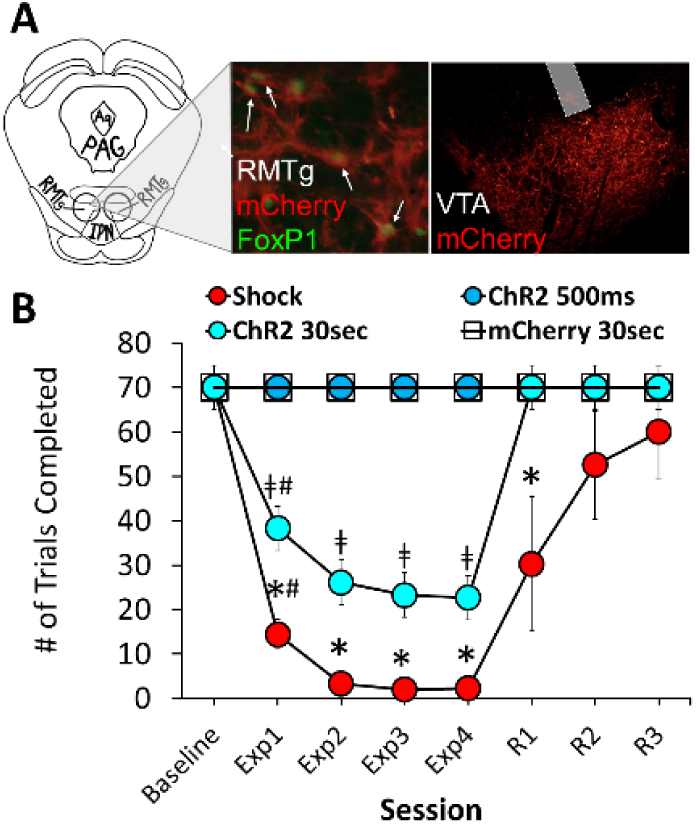
Optical stimulation of the RMTg◊VTA pathway suppressed lever pressing for food reward. **A)** Rats received bilateral injection of virus encoding channelrhodopsin-2 (ChR2) or control vector (mCherry) into the RMTg (left), and optical fibers were implanted above axon terminals in the VTA (right). **B)** Contingent footshock (0.5mA, 500ms) immediately after each completed trial significantly suppressed lever pressing for food reward. Comparable duration (500ms) optical stimulation of the RMTg-VTA pathway was ineffective at reducing food seeking, while sustained longer duration (30sec) optical stimulation robustly suppressed responding in an effect that became stronger with repeated testing (L1 vs L2-L4). Similar duration (30sec) light delivery in mCherry controls did not affect responding for food, indicating selective effects of ChR2 activation. n=5-8 per group; *p<0.05 shock vs all other groups; ╪p<0.05 ChR2 30sec vs all other groups; #p<0.05 within subjects effects of Exp1 vs Exp2-4.

### 3.2 Assessing effects of RMTg-VTA stimulation under conditions of expanded choice

Results from the Single Reward Task suggest that when rats are given limited response options (i.e. to “press or suppress” responding for food) RMTg-VTA stimulation is proximally effective to inhibit responding, yet food seeking rapidly resumed once the stimulation was terminated. Still, there were indications of more nuanced effects on learning (i.e. behavioral suppression was greater after repeated light-paired sessions in the ChR2 30sec stimulation group) that may have been masked during recovery sessions given the strong motivational drive in food-restricted animals. To therefore determine whether RMTg–VTA stimulation induces more enduring effects on learning when given alternative response options, we developed a Dual Reward Choice Task (see **Methods**) where responding for one of two distinctly-flavored food rewards was associated with pathway activation. In this task, we incorporated two complementary methods to stimulate RMTg efferents to the VTA, including chemogenetic conditioning and timing-specific optogenetic modulation. To determine if differences exist in baseline preference for the two food rewards (chocolate-or vanilla-flavored food pellets), all rats received three pre-stimulation choice sessions comprising 70 trials where they were permitted to freely respond for the two rewards. Choices for the different flavors averaged across the three pre-stimulation sessions showed ∼56% of rats chose vanilla more frequently, with ∼44% choosing chocolate more often (Figure 3A). Despite these individual differences in reward preference, there was no statistical difference in baseline intake of the two flavors irrespective of experimental manipulation (*F*(3,28) = .206, *p* = 0.891), and rats chose one of the two reward options approximately 65-70% of the time (Figure 3B). The magnitude of this baseline preference was consistent across experimental conditions and methodological approaches (i.e. chemogenetics vs optogenetics).

**Figure 3:**
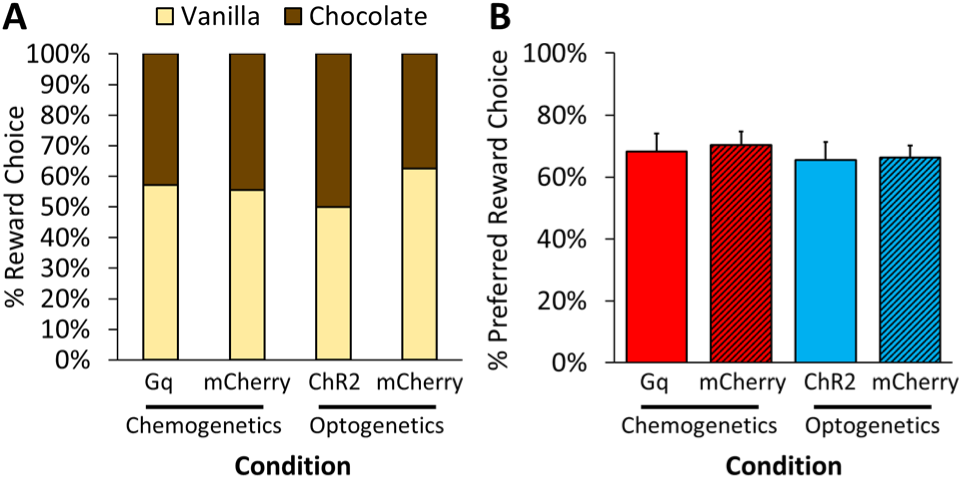
Initial flavor preference determined by three pre-stimulation choice sessions where rats were given the option to choose between vanilla-or chocolate-flavored food pellets. The flavor chosen more frequently averaged across the three sessions was deemed the “preferred” flavor. **A)** The proportion of choices by rats, separated by group, in the Dual Reward Choice Task. **B)** The proportion of preferred flavor chosen at baseline across the chemogenetic and optogenetic experiments. There were no significant differences across groups in the proportion of preferred flavor chosen at baseline (p>0.05).

### 3.3 Chemogenetic stimulation of the RMTg–VTA pathway conditions avoidance to the stimulation-paired reward

To test whether stimulation of the RMTg–VTA pathway is sufficient to induce lasting avoidance under expanded choice, we chemogenetically stimulated the RMTg–VTA pathway using a combination of retrogradely-transported cre recombinase in the VTA and cre-dependent excitatory Gq DREADDs in the RMTg (Figure 4A). Rats were then tested in the Dual Reward Choice Task where IP CNO injections were repeatedly paired with one of the two distinct food reward options, while repeated vehicle injections were paired with the other flavor (further described above in **Methods**). Given prior findings demonstrating RMTg–VTA stimulation causes real-time place aversion that becomes more robust with repeated testing [24], we hypothesized that stimulation of the RMTg–VTA pathway would condition avoidance against individual baseline choice biases, and this aversive learning effect would persist long after the initial conditioning.

**Figure 4:**
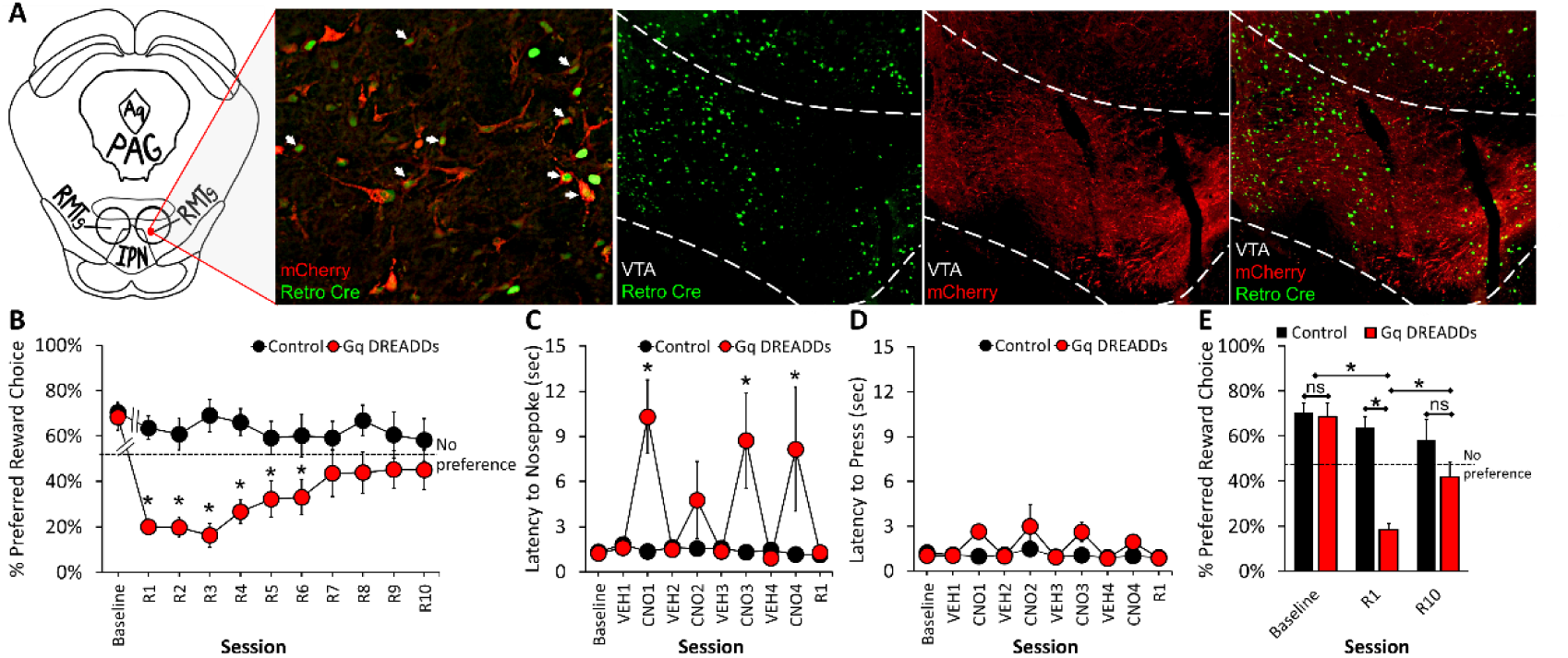
Pathway-specific chemogenetic stimulation of the RMTg–VTA pathway conditioned to the preferred food reward induces robust avoidance to the stimulation-paired reward. **A)** Rats received bilateral injections of cre-dependent excitatory Gq DREADDs or control vector (mCherry) in the RMTg (red) and bilateral injections of a retrograde cre-inducing virus in the VTA (green). **B)** Rats expressing Gq DREADDs exhibited a profound shift in preference away from the stimulation-conditioned reward option during post-stimulation “recovery” sessions (R1-R6). Line breaks between baseline and R1 represent conditioning sessions with CNO and vehicle. **C)** During conditioning sessions, CNO caused a significant increase in latency to initiate trials via nosepoke in Gq DREADDs rats, but **D)** no difference was observed in latency to press for reward. **E)** Gq DREADDs rats exhibited a robust conditioned avoidance to the CNO-paired reward in early choice testing (R1) that returned to baseline by R10. n=7-9 per group; *p<0.05 vs control (B, C, E) or R1 DREADDs vs R10 DREADDs (E).

Consistent with our predictions, Gq DREADDs rats showed a robust shift away from the initially preferred reward option (main effect of Group: *F*(1,14) = 14.146, *p* = 0.002; Figure 4B), indicating significant avoidance of the stimulation-paired reward. When examining reward preference across testing, we found a significant main effect of Session (*F*(3.123,43.721) = 7.904, *p* < 0.001) indicating the proportion of choices for the initially-preferred flavor shifted after conditioning with RMTg–VTA stimulation towards a robust and persistent avoidance of the stimulation-paired flavor. We also found a significant Session X Group interaction (*F*(3.123,43.721) = 6.560, *p* < 0.001) with post hoc analyses indicating the most profound change in preference occurred during earlier post-conditioning “recovery” sessions (R1-R6 *p* < 0.03; session 8 *p* = 0.008; DREADDs vs mCherry). While rats expressing Gq DREADDs exhibited a significant reduction in the proportion of responses towards the initially preferred reward option (within subjects R1-R5, *p* < 0.001 vs baseline), control rats (mCherry) were remarkably consistent in maintaining their initial preference across test sessions, indicating the effect of CNO on conditioned avoidance was not better explained by non-specific effects of the drug (p>0.05 vs mCherry baseline at all time points; Figure 4B).

Previous studies have shown RMTg lesions affect motor activity [17]. To therefore assess whether chemogenetic stimulation of the RMTg impacted performance during conditioning between baseline and R1 testing in the Dual Choice Task we examined latency to initiate trials via nosepoke in the food receptacle and latency to receive reward via pressing the central lever. For latency to initiate trials, Gq DREADDs rats took significantly longer to nosepoke during conditioning sessions compared to controls (main effect of Group: *F*(1,13) = 7.990, *p* = 0.014). These differences were driven by sessions where CNO was administered in rats expressing DREADDs (within subjects effect of Session: *F*(2.014,26.182) = 4.397, *p* = 0.022; Group X Session: *F*(2.014, 26.182) = 4.869, *p* .016; Figure 4C). Notably, when assessing latency to press the central lever following successful nosepoke, no significant differences were found between the control group and Gq DREADDs rats, nor were there significant within-subjects differences across sessions (p>0.05; Figure 4D), indicating that although RMTg–VTA stimulation made rats slightly slower to initiate trials during CNO conditioning, they were still able to very rapidly lever press to receive the reward. Importantly, all rats completed every trial in each conditioning session regardless of experimental condition ensuring equal exposure to the two flavors across groups (data not shown).

To more directly compare the proportion of choices towards the initially preferred flavor we assessed responses at baseline versus early (R1) and late (R10) testing, revealing a significant main effect of Group (*F*(1,14) = 10.843, *p* = 0.005), Session (*F*(1.233,17.268) = 14.178, *p* < .001), and a significant Group X Session interaction (*F*(1.233,17.268) = 8.174, *p* = 0.008). Post hoc analyses revealed a dramatic shift towards the non-preferred flavor at R1 in DREADDs rats (*p* < 0.001) (Figure 4E). No statistical differences were found between control and Gq DREADDs animals by R10 (*p* = 0.202), indicating that over the ∼2 weeks of post-conditioning testing rats expressing Gq DREADDs ultimately returned to their initial preference prior to conditioning.

### 3.4 Real-time optogenetic stimulation of the RMTg–VTA pathway induces long-lasting avoidance of the stimulation-paired reward

Building on our chemogenetic findings which demonstrated a robust, but transient, avoidance of the stimulation-conditioned flavor, we next determined whether timing-specific optogenetic RMTg–VTA activation produces real-time and persistent avoidance. To test this, we injected virus encoding ChR2 or control (mCherry) into the RMTg and implanted optical fibers above RMTg axon terminals in the VTA (Figure 5A). Rats were then tested in the same Dual Reward Choice Task described above; however, instead of chemogenetic conditioning rats underwent four light-paired sessions where each response for the initially-preferred reward at baseline was immediately followed by 30sec (33Hz) optical RMTg–VTA stimulation (to mimic the light parameters found to be effective in the Single Reward Task (Figure 2)).

**Figure 5:**
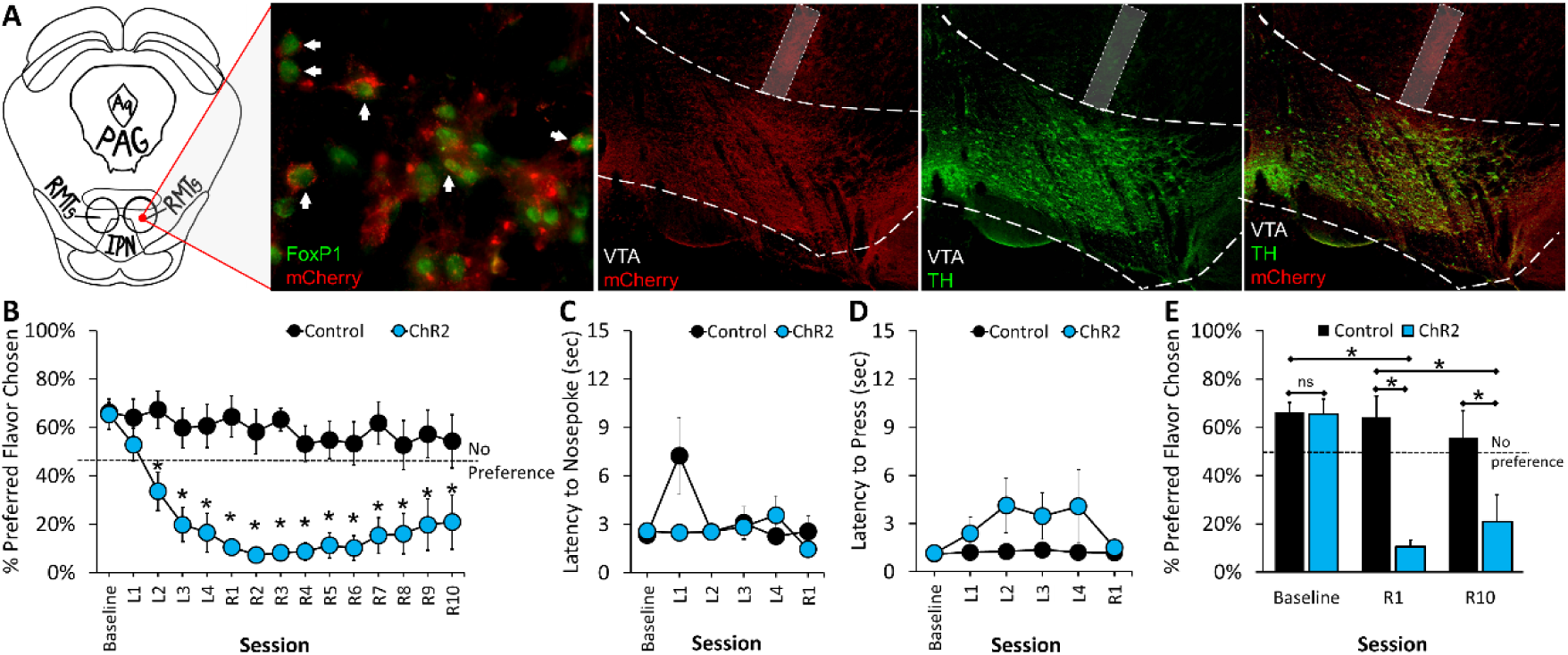
Temporally-specific optogenetic RMTg-VTA stimulation induces long-lasting avoidance when contingently paired to responding for the initially preferred reward option. **A)** Rats received bilateral injections of ChR2 or mCherry control in the RMTg (red) and optical fibers were implanted bilaterally above RMTg axons in the VTA (red). VTA sections were stained for tyrosine hydroxylase (TH) (green) to identify dopaminergic cells and FoxP1 (green) was used to delineate the RMTg. **B)** Rats expressing ChR2 exhibited a profound shift in preference away from the reward coupled to RMTg-VTA stimulation beginning in light session 2 (L2) and persisting throughout the remainder of testing. No differences were observed in either **(C)** latency to nosepoke to initiate trials or **(D)** latency to lever press for reward during light-paired sessions. **E)** Rats expressing ChR2 demonstrated enduring shifts in reward choice across both early (R1) and late (R10) testing in post-stimulation recovery sessions. n=8 per group; *p<0.05 vs control.

Rats expressing ChR2 robustly avoided the stimulation-paired reward option and shifted their choices toward the previously nonpreferred alternative (main effect of Group: *F*(1,14) = 27.579, *p* < 0.001; within subjects effects of Session: *F*(2.554,35.760) = 6.471, *p* = 0.002; Figure 5B). Beginning with the second session of real-time optogenetic RMTg-VTA stimulation, ChR2-expressing rats displayed a robust avoidance of the previously preferred flavor across light sessions 2 – 4 and across all 10 recovery sessions (all *p*-values < .05) when compared to controls (Session X Group interaction: *F*(2.554, 35.760) = 3.451, *p* = 0.033; Figure 5B). Notably, control rats did not alter their initial preference from baseline across any of the subsequent test sessions (all *p*-values = 1.000), indicating no non-selective effects of light delivery on behavior.

As mentioned above, modulation of the RMTg is known to affect locomotion. Unlike the chemogenetic approach, however, optogenetics allows for precise time-locked stimulation to the period immediately following the reward-seeking response (lever press), thereby minimizing overlapping influences of stimulation on earlier trial phases. In contrast to the effects of chemogenetic conditioning on latency to initiate trials, we found no significant differences in nosepoke latency during light-paired sessions (p>0.05; Figure 5C), nor were there any effects on lever press latency (*p*> 0.05; Figure 5D).

As with the chemogenetic experiment above, we compared the proportion of preferred flavor chosen at baseline, R1, and R10, to test for enduring effects of optogenetic RMTg-VTA stimulation. Analyses showed significant main effects of Group (*F*(1,14) = 16.007, *p* = 0.001) and Session (*F*(1.649,23.091) = 10.752, *p* < 0.001), and a significant Group X Session interaction (*F*(1.649,23.091) = 7.224, *p* = 0.006). Post hoc analyses revealed that ChR2 rats exhibited a significant reduction in the proportion of choices towards the initially preferred reward that persisted from R1 (*p* < 0.001) through R10 (*p* < 0.05) (Figure 5E).

### 3.5 RMTg–VTA circuit stimulation does not cause generalized reductions in effort-based responding or induce anhedonia

Previous studies have shown that inhibition of VTA DA neurons can produce depressive-like symptoms including anhedonia and reduced motivation to lever press for food [29, 30]. Therefore, to assess whether changes in reward choice caused by stimulation of the RMTg–VTA pathway extend to more generalized changes in motivation or affect, a separate group of rats expressing ChR2 or mCherry in the RMTg with optical fibers in the VTA were trained in the Dual Reward Choice Task described above. Rats received 4 light-paired stimulation sessions to induce real-time avoidance of the stimulation-paired flavor, and the following day they were assessed in 8, twice-daily, progressive ratio sessions, separately testing effort-based responding for each reward type after the stimulation. This allowed us to determine enduring effects of the prior stimulation on motivation to obtain either of the two reward options at a time that coincided with the long-lasting avoidance observed in prior testing when both reward options were available.

Replicating our chemogenetic findings, ChR2 rats exhibited significant avoidance of the stimulation-paired reward and a shift in preference away from the stimulation-paired flavor (main effect of Group: *F*(1,14) = 13.007, *p* = 0.003; Session X Group interaction *F*(1.831,25.641) = 12.534, *p* < 0.001; Figure 6A), with ChR2 rats shifting preference away from the stimulation-paired reward across light sessions 2-4 (*p* < 0.05 vs control). Notably, when subsequently tested under the progressive ratio schedule of reinforcement, we found no difference in average PR breakpoint between ChR2 rats and controls (between subjects effect of Group: *F*(1,14) = 0.001, *p* = 0.979; Figure 6B). Within subjects tests of reward type (light-paired versus no light), however, showed that all rats, irrespective of group, responded more for the initially preferred (light-paired) flavor versus the non-preferred reward (*F*(1,14) = 7.679, *p* = 0.015), indicating that even though rats will avoid the light-paired flavor when given a choice, they will still equally work for it if no alternative is provided. No significant interaction effects in PR breakpoint were observed (*p* = 0.963).

**Figure 6:**
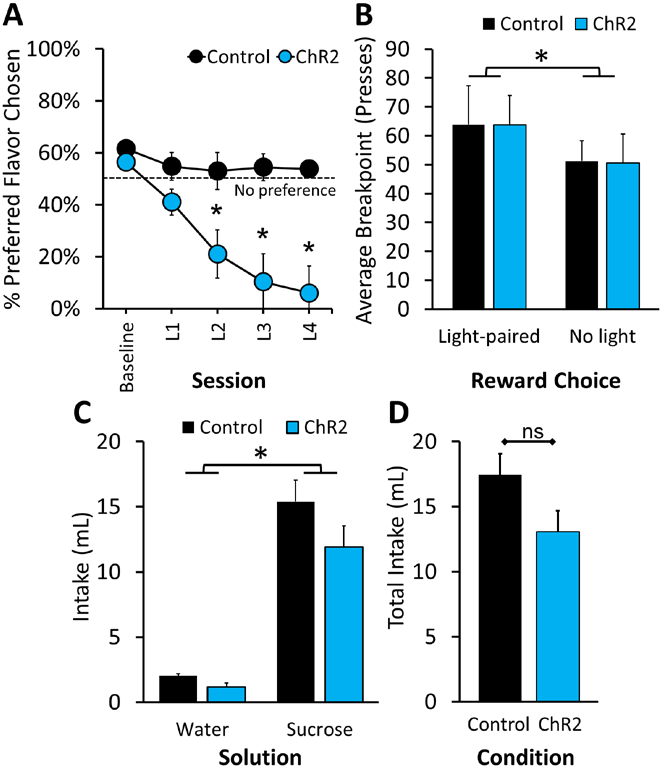
Optogenetic stimulation of the RMTg–VTA pathway does not alter effort-based responding for individual rewards or induce general reductions in reward processing. **A**) We replicated previous testing demonstrating contingent RMTg–VTA stimulation caused a robust shift in reward preference in the Dual Reward Choice Task. **B)** Immediately after testing for reward preference, the same animals underwent progressive ratio (PR) testing for each individual flavor in twice-daily testing where only a single reward was available. There was no difference in PR breakpoint for either flavor in ChR2 or control rats, although rats responded more for the initially-preferred flavor irrespective of virus condition. After PR testing rats were removed from food restriction and tested for sucrose preference. ChR2 and control rats exhibited similar preference for sucrose over water **(C)**, but no differences in intake of the individual solutions or **(D)** total intake, were observed. n=7-8 per group; *p<0.05 ChR2 vs control (A), light-paired vs no light (B), sucrose vs water (C).

After completing PR testing, rats were again given *ad libitum* food access and the next day began testing in a two-bottle sucrose preference task to determine if previous RMTg–VTA circuit stimulation caused long-lasting changes in reward processing. We found no significant effect of virus condition (ChR2 vs control; *p* = 0.083), but there was a significant within subjects effect of solution (sucrose vs water; *F*(1,12) = 101.757, *p* < 0.001) indicating robust preference for sucrose solution regardless of group (Figure 6C), while no significant interaction effect was found (*p* = 0.290). Similarly, no difference in total fluid intake was observed when combining water and sucrose consumption between ChR2 and control rats (*p* = 0.083; Figure 6D).

## 4.0 Discussion

Optimizing cost-benefit decision-making is fundamental for adaptive behavior, allowing organisms to maximize rewards while minimizing negative consequences. Disruptions in this process, however, are characteristic of several psychiatric disorders including mania and substance use disorder, where maladaptive reward seeking persists despite incurring often substantial costs [1, 31]. Greater understanding of the neural circuits that mediate avoidance learning to moderate reward seeking is therefore critical to identify novel targets for future treatment of compulsive disorders. Here, we demonstrate that stimulation of the RMTg–VTA circuit serves as an aversive teaching signal to modulate reward-seeking and situationally-dependent behavior. Specifically, activation of the RMTg–VTA pathway induces passive avoidance (behavioral inhibition) when faced with limited response options (i.e. lever press to receive reward and incur stimulation “cost” or suppress responding to avoid stimulation but miss out on reward). In situations where reward availability is more flexible (as in the Dual Choice Task), the RMTg–VTA circuit appears to support a different strategy oriented towards active avoidance, as pathway stimulation promotes shifts in subjective choice by deterring particular responses rather than non-selective behavioral suppression. Moreover, even though RMTg–VTA stimulation caused a dramatic shift in choice with subjects avoiding the stimulation-paired reward for many days after the initial learning, it did not produce generalized reductions in effort-based responding (progressive ratio) or anhedonia, as would be expected from non-selective and enduring changes in dopaminergic activity.

The multifaceted role of the RMTg revealed in the current studies is further supported by several key findings. For example, lesions of the RMTg or pathway-specific RMTg-VTA optogenetic inhibition profoundly impairs the ability to withhold responding for food reward in the context of impending footshock [18, 32], suggesting deficits in behavioral inhibition after inactivation of this circuit. It is perhaps not surprising then that experimental stimulation of this region suppresses behavioral responding, as demonstrated here. Given the recognized motoric effects of RMTg manipulations [17, 18], it is difficult to disentangle motivational effects of stimulating this circuit from the potential overlapping effects of changes in movement. The finding that lever pressing for reward rapidly resumed when stimulation was removed during “recovery” sessions in the Single Reward Task would seem to support this notion. It is only when an additional response option is made available in the Dual Reward Choice Task that we reveal rats are indeed still capable of volitional movement during (chemogenetic) or after (optogenetic) RMTg stimulation, albeit at a slightly slower rate (chemogenetics), and they will learn to avoid it if given the choice. Indeed, when given no other option for food reward rats appear able to overcome the presumed aversive response to the stimulation-paired flavor, as evidenced by the rapid resumption of lever pressing during recovery in the Single Reward Task and the lack of change in progressive ratio despite the observed shift in subjective choice in the Dual Reward Task. Yet when given the opportunity to avoid, we found a shift in preference that lasted for nearly two weeks after the initial stimulation, and likely longer if testing was more protracted, reflecting the active avoidance responses reported in prior place conditioning paradigms [19, 24, 33].

It is notable that with our chemogenetic approach, the conditioning strategy resembles a mix of Pavlovian and instrumental (operant) conditioning [34]. For example, CNO-induced activation of the RMTg–VTA pathway acts as a Pavlovian unconditioned stimulus—by not requiring behaviorally-contingent lever pressing to elicit its activation, as was the case for our optogenetic approach. Rather, activation of the pathway occurs regardless of their behavior, but in conjunction with motivated lever pressing for the preferred flavor, illustrating the operant component. Therefore, by allowing access to only one flavor option during any given conditioning session, we presumably induced a learned association between RMTg–VTA pathway activation and the action of lever pressing for the preferred flavor as well as to the preferred flavor itself. This is an important consideration given our findings in the Gq DREADDs group demonstrating a more transient avoidance response to the preferred flavor compared to timing-specific optogenetic manipulation. Given that RMTg–VTA stimulation occurred throughout the entire session, other associations with the experimental context alongside pressing for the preferred flavor may have occurred, potentially obscuring conditioning. Indeed, classic studies in associative learning have shown that irrelevant stimuli unrelated to the conditioning paradigm can disrupt the learned association between a response and its consequence [35].

The largest avoidance response (∼75% reduction) was seen in the recovery choice sessions immediately following chemogenetic conditioning suggesting that, over time, with access to both flavor options, the strength of the conditioned avoidance degrades. This extinction effect is commonly seen in classic fear conditioning paradigms where removal of the aversive (unconditioned) stimulus leads to extinction of the conditioned response [36], which in this case would be avoidance of the preferred flavor. Precisely timing optogenetic stimulation of the RMTg–VTA pathway as an immediate consequence for seeking out the rat’s preferred flavor produced a more robust and longer lasting avoidance response than what was seen in our chemogenetic experiment. This corroborates previous fear conditioning studies which have shown that continuity between the aversive stimulus, in this case RMTg–VTA stimulation, and the neutral stimulus (preferred flavor) has a substantial impact on the strength of the learned association as well as the length of time before the learned association is extinguished [37].

The use of two distinctly-flavored pellets to model different reward options may have important considerations for impacting gustatory and taste-related circuits in the brain including gustatory cortex, which encompasses parts of the insula and frontal operculum, as well as brainstem regions including the parabrachial nucleus (PBN) [38, 39]. Studies have shown that the PBN projects to the RMTg [40], and the PBN has also been shown to be recruited in assigning hedonic or aversive value to tastes [41]. Thus, inducing avoidance toward a particular reward option may recruit the PBN by perhaps altering the palatability of the food itself. Given the pathway-specific strategies used here, and the lack of change in effort-based responding for the reward previously paired with RMTg-VTA stimulation, this interpretation seems less likely but would be an important consideration of future work.

Given that avoidance in the Dual Choice Task persisted for many days after stimulation ceased, it suggests a learned association between the neural experience of RMTg–VTA stimulation and the behavioral response (i.e. nosepoke or lever press), the reward type, or both. Such associations are likely the result of changes in neuronal plasticity in the brain resulting from RMTg–VTA stimulation. Given RMTg efferents to the VTA predominantly synapse onto DA neurons [15–17], this would be a likely site of integration, although DA activity does not appear to be generally downregulated (given results of PR and SPT), and the RMTg has not been shown to undergo traditional forms of synaptic plasticity [42, 43]. Nevertheless, this does not rule out long-lasting changes in signaling between the RMTg or other inputs to the VTA, in DAergic or non-DAergic cells, or other potential downstream forebrain targets that may be influenced by RMTg-induced reductions in VTA activity, such as the nucleus accumbens or prefrontal cortex [44].

The present studies demonstrate that the RMTg, through its connections with the dopaminergic VTA, is sufficient to drive context-dependent avoidance behavior. Further, repeated contingent stimulation of this circuitry is capable of producing enduring changes in decision-making under conditions of expanded choice, in a manner that is selective to learned associations without inducing broad changes in reward processing which may otherwise drive depression, anxiety, and substance use disorder [45, 46]. These results provide key insights into the neural mechanisms that selectively drive long-lasting changes in behavioral responding for rewards, and future work will be critical to identify novel targets for the treatment of disorders ranging from inappropriately excessive avoidance (i.e. anxiety-related disorders and PTSD) to insufficient inhibitory control and persistent reward seeking, including substance abuse and behavioral addictions.

## Acknowledgements

The authors would like to thank Murphy Miller and Anna Caroline Toburen for assistance with behavioral studies and general research support. We would also like to thank Dr. Karl Deisseroth and the UNC Vector Core for making available to us viral constructs encoding channelrhodopsin-2 (AAV2-hSyn-hChR2(H134R)-mCherry) and control vector (AAV2-hsyn-mCherry), and the NIDA drug supply for sharing clozapine-N-oxide.

## Author contributions

J.R.W., M.D.L., and P.J.V. were involved in the conceptualization, methodology, and data collection of this research. E.L.C. assisted with data collection. J.R.W. and P.J.V. were responsible for data analysis, writing, and editing. M.D.L. and E.L.C. assisted with reviewing and editing the manuscript.

